# Acetylation of nuclear localization signal controls importin-mediated nuclear transport of Ku70

**DOI:** 10.1101/403485

**Authors:** H Fujimoto, T Ikuta, A Koike, M Koike

## Abstract

Ku70 participates in various intra-and extra-nucleic processes. For multifunctional control, machinery that precisely regulates the intracellular localization of Ku70 is essential. Recently, it was reported that acetylation of Ku70 regulates its function. Here, we demonstrate that specific lysine residues in Ku70 that are targets of acetylation are critical for regulating nuclear transport *in vivo*. Ku70-GFP fusion proteins transiently expressed in cultured cells localized in the nucleus, whereas mimicking acetylation of K553 or K556 in the Ku70 nuclear localization signal (NLS) by substituting these lysine residues with glutamine markedly decreased the nuclear localization of Ku70. Moreover, the Ku70-importin interaction was suppressed in the K553Q and K556Q mutants. Theoretical estimations indicated that the binding energy between the Ku70 NLS and importin-α decreases with acetylation of lysine residues in the Ku70 NLS, similar to the case when these lysine residues are substituted with glutamine. These results suggest that acetylation of specific lysine residues in the Ku70 NLS is a key switch that controls the localization of Ku70 by modulating interactions between Ku70 and nuclear transport factors.

## Introduction

Ku is a multifunctional protein originally identified as an autoantigen in the serum of patients with the autoimmune disease dipolymyositis-scleroderma overlap syndrome (Mimori et al, 1981). Later studies revealed that Ku functions in various cellular processes. For example, intranuclear Ku is involved in DNA repair, immunoglobulin gene rearrangement, telomere maintenance, and transcriptional regulation (Downs & Jackson, 2004; Featherstone & Jackson, 1999; Tuteja & Tuteja, 2000), whereas Ku expressed on the cell surface plays roles in cellular adhesion and proteolytic processes by interacting with matrix metalloproteinase 9 (Koike, 2002; Monferran et al, 2004; Paupert et al, 2007). Moreover, cytosolic or nuclear Ku binds to exogenous DNAs or proteins of viruses such as HIV, HTLV-1, adenovirus, HSV-2, and HVB (Anisenko et al, 2017; Frost et al, 2017; Li et al, 2016; Rushing et al, 2018; Sui et al, 2017; Wang et al, 2017), suggesting that Ku70 acts as a sensor during virus infection or as a host factor against some viruses. Among its many functions, Ku has been well characterized as a DNA double-strand-break (DSB) sensor in the process of non-homologous end-joining (NHEJ), which is a major pathway for the repair of DSBs arising from exposure to ionizing radiation or chemical agents in mammalian cells. After Ku recognizes and binds DSB sites in DNA, it provides a foothold for the formation of a complex involving NHEJ-related components such as DNA-PKcs, XRCC4, XLF, and ligase IV (Koike & Koike, 2008; Koike et al, 2011; Mahaney et al, 2009).

Ku70 is a subunit of Ku that associates with its homologous protein, Ku80, to form Ku heterodimers in the nucleus. Interestingly, the cytoplasmic distribution of Ku70 was found to be significantly higher than that of Ku80 (Cohen et al, 2004a), suggesting that Ku70 and Ku80 function separately in the cytoplasm. For instance, cytoplasmic Ku70 independent of Ku80 suppresses apoptosis by sequestering the pro-apoptotic factor Bax from the mitochondria (Cohen et al, 2004a; Subramanian et al, 2005). Ku70 also binds to and stabilizes FLIP by suppressing its polyubiquitination. Inhibition of histone deacetylase 6 (HDAC6) activity promotes the acetylation of Ku70 and disrupts the Ku70/FLIP interaction, resulting in induction of apoptosis by caspase 8 (Kerr et al, 2012). Furthermore, it was reported that binding of p18-cyclin E with cytosolic Ku70 leads to the dissociation of Bax from Ku70, with subsequent activation of Bax and induction of mitochondria-mediated apoptosis (Mazumder et al, 2007). Despite the polyfunctionality of Ku70 in the nucleus as a heterodimer with Ku80 and in the cytoplasm as a single protein, the mechanism of Ku70 translocation from the cytoplasm to the nucleus is poorly understood.

Previously, we determined that the nuclear localization signal (NLS) of human Ku70 is located in the region between residues K539 and K556 and that the NLS is bound and transported to the nuclear rim by the nuclear transport factors importin-α/importin-β complexes (Impα/Impβ) (Koike et al, 1999c). The molecular structure of complexes composed of the Ku70 NLS and Impα was solved by X-ray crystallographic analyses (Takeda et al, 2011). The formation of a complex between Ku70 and heat shock cognate protein 70 was shown to prevent the NLS of Ku70 from accessing Impα, thus inhibiting the nuclear translocation of Ku70 in AR42J rat pancreatic acinar cells (Lim et al, 2008). These findings indicated that Ku70 is translocated to the nucleus via the classical Impα/Impβ-mediated nuclear import pathway. In addition, the Ku70 NLS contains five lysines (K539, K542, K544, K553, and K556) that were identified as targets for acetylation *in vivo* (Cohen et al, 2004a; Koike et al, 1999c). Acetylation is a major protein posttranslational modification (PTM) that is thought to promote conformational changes that control association with or dissociation from other biomolecules by neutralizing positive charges (Yang & Seto, 2008). Thus, it is hypothesized that acetylation of the five lysine residues in the Ku70 NLS acts as a switch controlling the translocation of Ku70 from the cytoplasm to the nucleus.

Here, we report the effects of lysine acetylation in the NLS of Ku70 using both experimental and theoretical approaches and demonstrate that recombinant Ku70 mutants in which specific NLS lysine residues were substituted with glutamine to mimic acetylation exhibited reduced translocation to the nucleus. These findings suggest that translocation of Ku70 to the nucleus is controlled by the acetylation of specific lysine residues in the NLS.

## Results

### Acetylation-mimicking mutation of lysine residues 553 and 556 but not lysine residues 539, 542, or 544 inhibit the nuclear localization of Ku70

Amino acid sequences in the putative NLS are highly conserved among primates. In particular, five lysine residues (K539, K542, K544, K553, and K556) in human Ku70, which are targets for acetylation, are highly conserved among mammalian and avian Ku70 homologues, although K544 is replaced with a similar basic residue, arginine, in canine species (Koike et al, 1999c; Koike et al, 2017a) (Figure 1).

**Figure 1.**
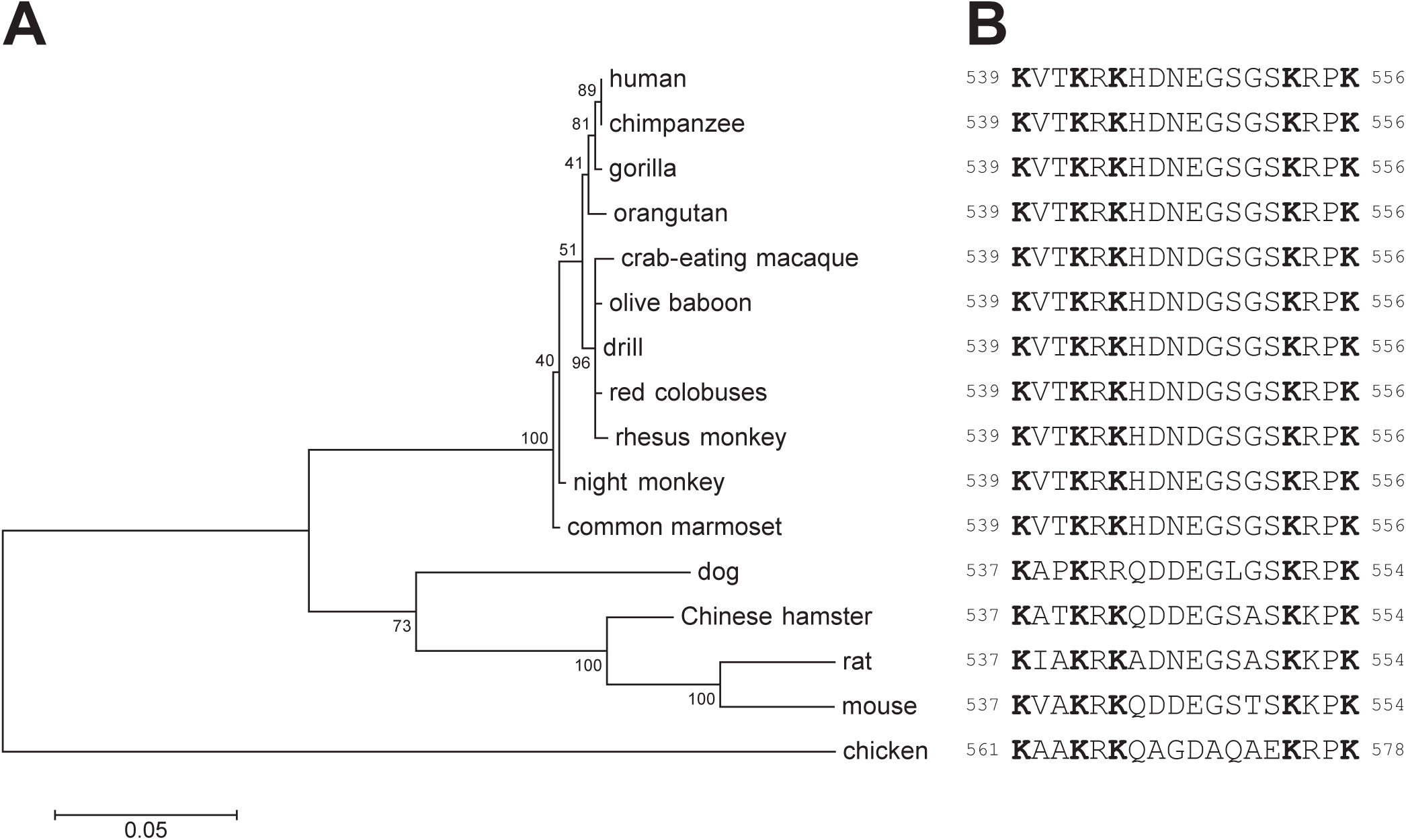
Acetylation sites in the NLS of human Ku70 are conserved in the putative NLS among the primates. **A** Phylogenetic tree for Ku70 amino acid sequences constructed by the maximum-likelihood method using MEGA 5.2.2 software (Tamura et al, 2011). The JTT-f model (Adachi & Hasegawa, 1996; Jones et al, 1992) was applied as a substitution model to estimate evolutionary distances between amino acid sequences. All positions containing gaps were eliminated. Bootstrap tests were replicated 1000 times for each node. The NCBI Reference number of the amino acid sequences used for the analysis are as follows: human (*Homo sapiens*): NP_001460.1, chimpanzee (*Pan troglodytes*): NP_001267434.1, gorilla (*Gorilla gorilla gorilla*): XP_004063587.1, orangutan (*Pongo abelii*): NP_001126888.1, crab-eating macaque (*Macaca fascicularis*): BAE00743.1, olive baboon (*Papio anubis*): XP_009215653.1, drill (*Mandrillus leucophaeus*): XP_011834155.1, red colobus (*Piliocolobus tephrosceles*): XP_023077638.1, rhesus monkey (*Macaca mulatta*): EHH20277.1, night monkey (*Aotus nancymaae*): XP_021522829.1, common marmoset (*Callithrix jacchus*): XP_008989116.1, dog (*Canis lupus*): LC195221, Chinese hamster (*Cricetulus griseus*): XP_007618650.1, rat (*Rattus norvegicus*): NP_620780.2, mouse (*Mus musculus*): NP_034377.2, and chicken (*Gallus gallus*): NP_990258.2. All amino acid sequences of Ku70 were aligned using Clustal W software, version 2.1 (Larkin et al, 2007). **B** Alignment of amino acid sequence of the putative Ku70 NLS of each organism listed in panel **A**. Bold letters indicate acetylation sites deduced from those identified in human Ku70 NLS by Cohen *et al.* (2004). The numbers on either side of the NLS indicate residue numbers for each organism.

To examine whether mutations at acetylation sites in the Ku70 NLS affect its nuclear localization, we carried out site-directed mutagenesis experiments in which Ku70 was labeled with EGFP and each lysine in the Ku70 NLS was substituted with an uncharged glutamine (KQ mutants mimicking acetylated lysine), or a charge-conservative arginine (KR mutants mimicking non-acetylated [deacetylated] lysine) residue (Figure 2).

**Figure 2.**
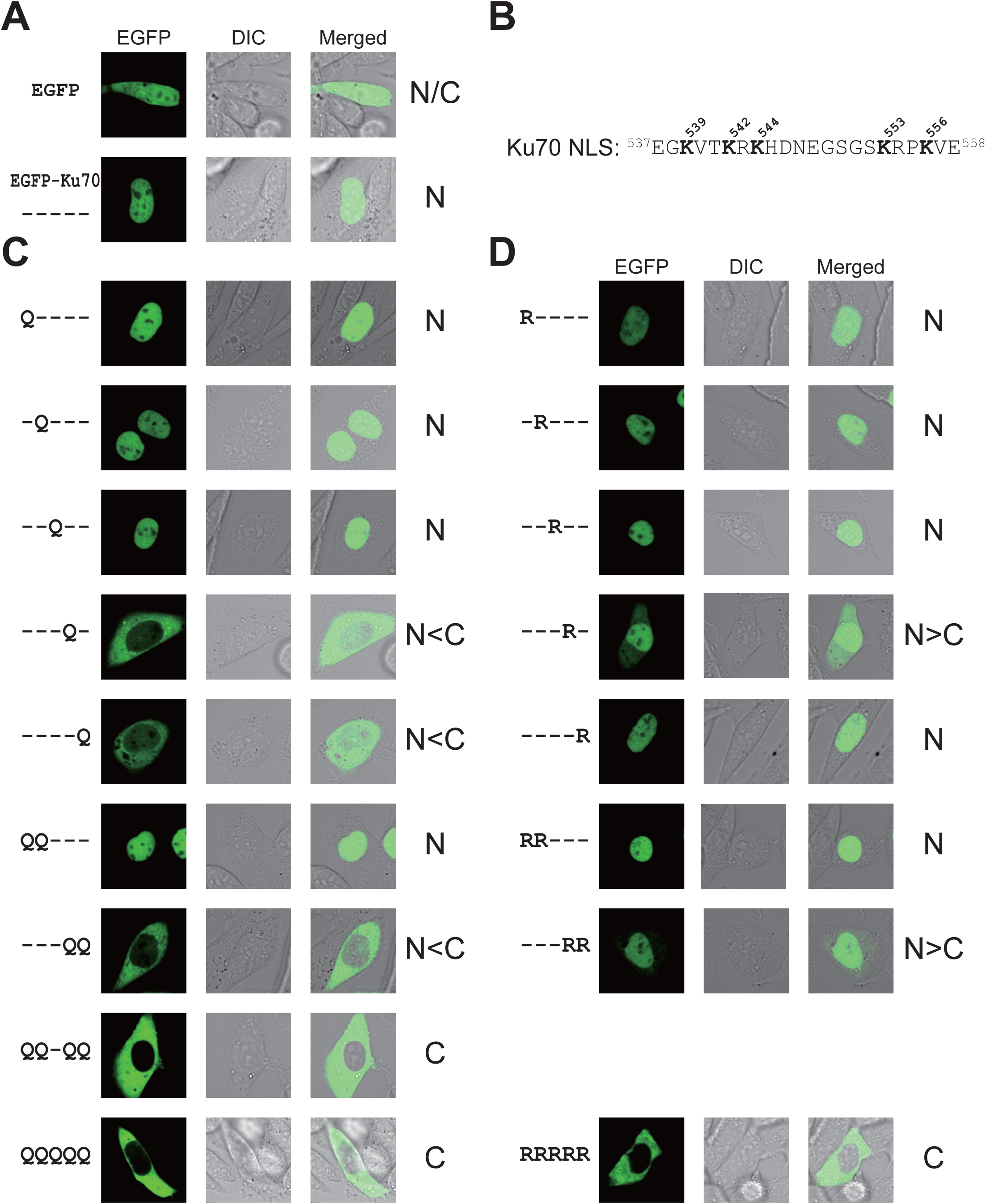
Nuclear localization activity of Ku70. **A** EGFP signal, differential interference contrast (DIC), and merged images of the EGFP–wild-type Ku70 fusion protein and EGFP only in xrs-6 cells. **B** Amino acid sequence of the Ku70 NLS. Five lysine (K) residues that were reported as targets of acetylation are indicated in bold letters. Numbers on either side of the sequence and above the lysine residues indicate residue positions in Ku70. **C, D** EGFP signal of acetylation-mimicking fusion proteins (KQ mutants) and deacetylation-mimicking fusion proteins (KR mutants) in xrs-6 cells. Data information: In **A**, **C**, and **D**, positions of substitution among the five lysine residues are indicated to the left of each panel; for example, “Q----” indicates a single mutation, K539Q. The predominance of localization in xrs-6 cells exhibiting translocation of the EGFP signal in each case is indicated to the right (N: nucleus, C: cytoplasm).

Translocation of Ku70 from the cytoplasm to the nucleus was almost completely abolished when all five lysine residues or four lysine residues (except K544) were replaced with glutamine residues (QQQQQ or QQ-QQ in Figure 2C). The double mutant EGFP-Ku70 K553Q/K556Q (---QQ in Figure 2C) was detected mostly in the cytoplasm, but the double mutant EGFP-Ku70 K539Q/K542Q (QQ--- in Figure 2C) was detected primarily in the nucleus. Mutants in which all five lysine residues were replaced with alanine residues (KA mutants) exhibited similar localization of EGFP-Ku70 (Figure EV1). These results suggest that inhibition of the translocation of Ku70 from the cytoplasm to the nucleus is greater when a substituted lysine residue is located closer to the C-terminus of the Ku70 NLS. Single mutations at each acetylation site support this presumption. EGFP-Ku70 K553Q (---Q-in Figure 2C) or EGFP-Ku70 K556Q (----Q in Figure 2C) were detected primarily in the cytoplasm, whereas the wild-type Ku70 fusion proteins EGFP-Ku70 (----- in Figure 2A), EGFP-Ku70 K539Q (Q---- in Figure 2C), EGFP-Ku70 K542Q (-Q--- in Figure 2C), or EGFP-Ku70K544Q (--Q -- in Figure 2C) were detected primarily in the nucleus, suggesting that two of the five lysine residues in the Ku70 NLS (K553 and K556) play more important roles in nuclear import of Ku70. In contrast, the inhibition of Ku70 translocation to the nucleus was not significant when K539, K542, K544, K553, or K556 were replaced separately or when K539/K542 or K553/K556 were substituted concurrently with arginine residues (Figure 2D), although translocation of Ku70 to the nucleus was almost completely abolished when all five lysine residues were substituted with arginine residues (RRRRR in Figure 2D).

These findings indicate that the substitution of a glutamine residue for K553 or K556 affects the ability of Ku70 to localize to the nucleus, suggesting that Ku70 likely remains in the cytoplasm when K553 and/or K556 in the NLS are acetylated and translocates to the nucleus when K553 and/or K556 are not acetylated.

### Mimicking acetylation of K539 or K542 has no effect on accumulation of Ku70 at sites of DNA damage or heterodimerization with Ku80 in living cells

Mimicking acetylation of either or both lysine residues (K539 and K542) had a negligible impact on nuclear transport (Figure 2C), although substitution of these two lysine residues with alanine residues partially inhibited nuclear transport in some cells (Figure EV1B). It was reported that the substitution of either K539 or K542 with glutamine abolished the ability of Ku70 to bind Bax (Cohen et al, 2004a); therefore, acetylation of these two lysine residues is thought to facilitate Bax-mediated apoptosis. Chen *et al*. also demonstrated that the same KQ mutants at K539 or K542 suppressed the DNA end-binding activity of Ku70 *in vitro* (Chen et al, 2007).

We examined the effect of the K539Q and/or K542Q mutations on Ku70 accumulation at DSBs *in vivo*. As Ku80 is essential for Ku70 accumulation in xrs-6 cells at DSB sites induced by irradiation with a 405-nm laser (Koike & Koike, 2008), the subcellular localization and dimerization of EGFP-Ku70 and its mutants (EGFP-Ku70 K539Q, EGFP-Ku70 K542Q, or EGFP-Ku70 K539Q/K542Q) were examined in xrs-6 cells co-transfected with ECFP-Ku80, using EGFP tags and the specific antibody 162, which recognizes Ku heterodimers but not Ku70 or Ku80 monomers (Wang et al, 1994). As shown in Figure 3A, when ECFP-Ku80 was co-expressed with EGFP-Ku70, as well as the double KR mutant, EGFP-Ku70 K539R/K542R, antibody 162 readily detected the Ku heterodimer in the nucleus. When ECFP-Ku80 was co-expressed with EGFP, the Ku heterodimer was not detected (Figure 3A). Antibody 162 detected the Ku heterodimer in the nucleus of all cells examined, even when EGFP-Ku70 K539Q, EGFP-Ku70 K542Q, or EGFP-Ku70 K539Q/K542Q and ECFP-Ku80 were expressed together, indicating that heterodimerization of Ku70 and Ku80 in xrs-6 cells was not inhibited by these substitutions. To examine whether these mutants accumulate at DSBs, we transiently co-expressed each EGFP-Ku70 or its mutants with ECFP-Ku80 in xrs-6 cells and microirradiated the cells using a 405-nm laser. Both EGFP-Ku70 (wild type) and EGFP-Ku70 K539R/K542R accumulated, but EGFP alone did not accumulate after laser microirradiation (Figure 3B). EGFP-Ku70 accumulated at microirradiation sites in living cells even when either or both K539 and K542 were substituted with glutamine residues (K539Q, K542Q, K539Q/K542Q).

**Figure 3.**
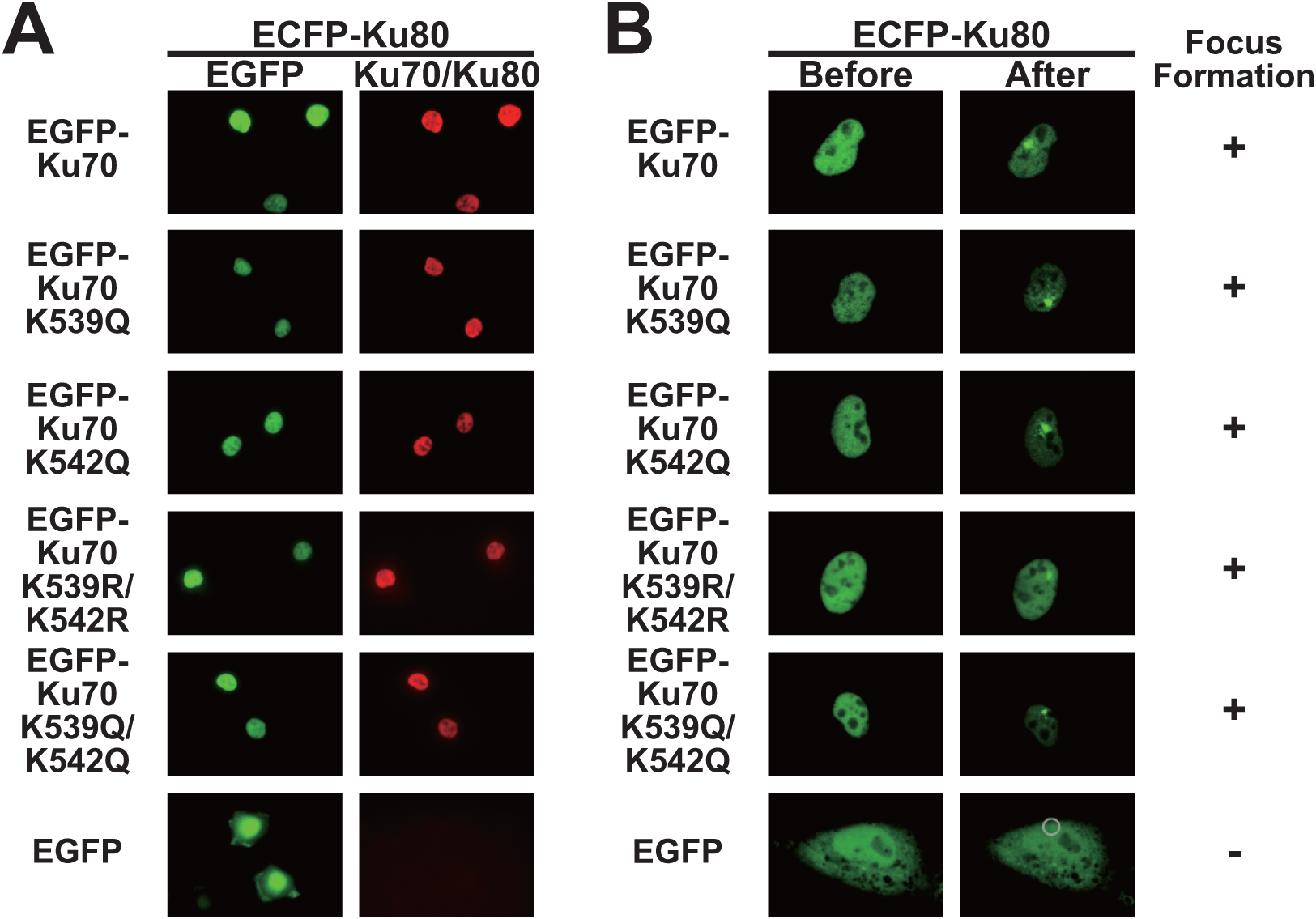
Substitution of Ku70 lysine residues 539 and 542 with acetylation-mimicking mutations do not diminish heterodimer formation activity with Ku80 in the nucleus or accumulation at DNA damage sites *in vivo*. **A** Direct interaction between EGFP-Ku70 or its mutant and ECFP-Ku80 *in vivo.* Ku80-deficient xrs-6 cells were transiently cotransfected with pECFP-Ku80 and pEGFP-Ku70, pEGFP-Ku70 K539Q, pEGFP-Ku70 K542Q, pEGFP-Ku70 K539R/K542R, pEGFP-Ku70 K539Q/K542Q, or pEGFP. The cells were fixed and stained with Ku70/Ku80 antibody 162 specific for the Ku70/Ku80 complex. The subcellular localization of EGFP-tagged proteins (green) and Ku70/Ku80 complex (red) was examined in the same cells by confocal laser scanning microscopy. **B** xrs-6 cells were transiently cotransfected with pECFP-Ku80 and pEGFP-Ku70, pEGFP-Ku70 K539Q, pEGFP-Ku70 K542Q, pEGFP-Ku70 K539R/K542R, pEGFP-Ku70 K539Q/K542Q, or pEGFP. The accumulation of EGFP-tagged protein at sites of DNA damage induced by irradiation with a 405-nm laser was examined. Live-cell images were taken before and 60 s after microirradiation. Accumulation (+, accumulated; -, not accumulated) at DNA damage sites is summarized to the right. Gray circle in the right image of EGFP denotes the site of microirradiation.

These findings suggest that the substitutions of any of two lysine residues (i.e., K539 and K542) in the acetylation site of Ku70 has minimal effect on nuclear localization and accumulation at DSB sites, although binding activity of Ku heterodimers to DSB sites is reportedly decreased *in vitro* if K539 or K542 is substituted with glutamine.

### Substitution of Ku70 lysine residues K553 and K556 to mimic acetylation diminishes its interaction with importin-α□importin-β

To confirm the significance of K553 and K556 in the Ku70 NLS for Ku70 translocation to the nucleus, we performed a microinjection assay using the GST-NLS-GFP protein system. As we reported previously (Koike et al, 1999c), the purified GST-Ku70 (539-556)-GFP fusion protein exhibited efficient nuclear translocation within 30 min of incubation at 37°C when GST-Ku70 (539-556)-GFP was microinjected into the cytoplasm of HeLa cells (Figure 4A). The positive control, GST-NLSc-GFP, also exhibited efficient nuclear localization, whereas the negative control, GST-GFP, exhibited no nuclear localization. The Ku70 NLS quadruple mutant, GST-Ku70 (539-556, K539Q/K542Q/K553Q/K556Q)-GFP, and double mutant, GST-Ku70 (539-556, K553Q/K556Q)-GFP, lost nuclear translocation activity, indicating that mimicking the acetylated state of two specific lysine residues, K553 and K556, reduces the activity of the Ku70 NLS (Figure 4A).

**Figure 4.**
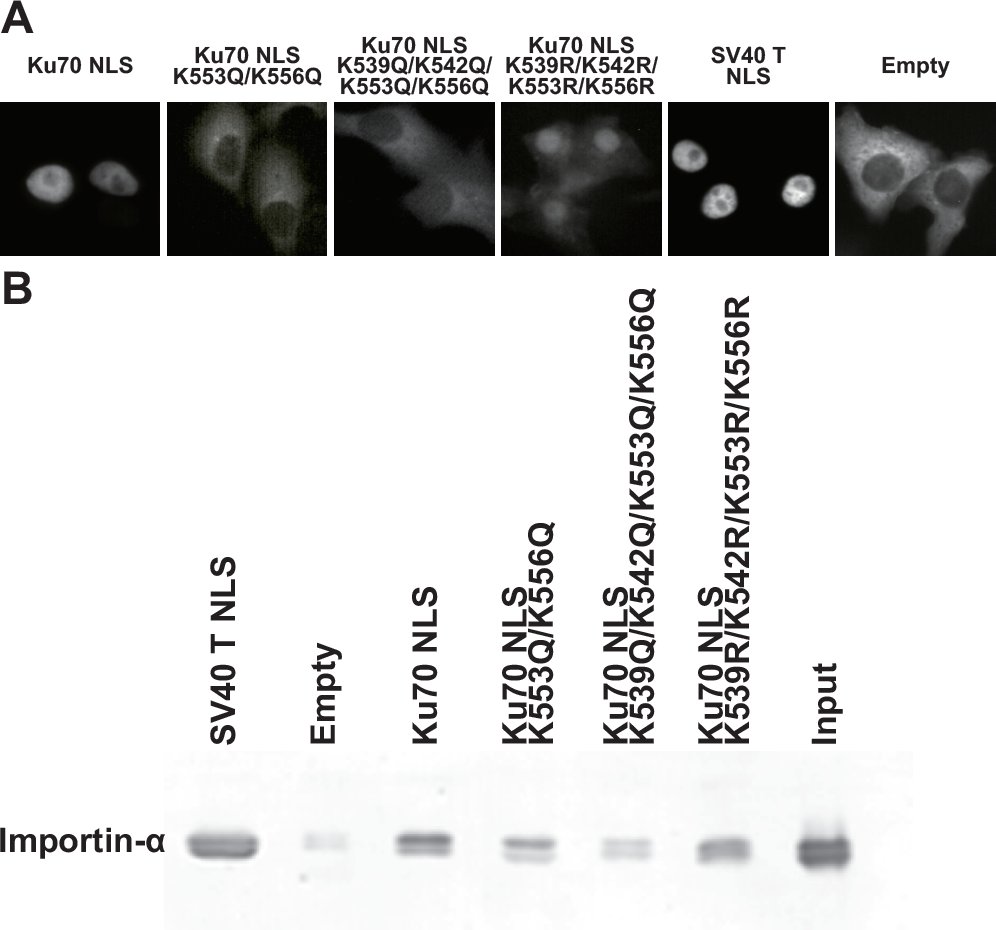
Effect of substitution of lysine residues in the Ku70 NLS with acetylation-mimicking mutations on nuclear entry and binding to Impα. **A** Microinjection of HeLa cells with recombinant GST-Ku70(539-556)-GFP and its mutant proteins. Affinity-purified recombinant proteins, GST-Ku70(539-556)-GFP (Ku70 NLS), GST-Ku70(539-556,K553Q/K556Q)-GFP(Ku70NLSK553Q/K556Q),GST-Ku70(539-556,K539Q/K542Q/K553Q/K556Q)-GFP (Ku70 NLS K539Q/K542Q/K553Q/K556Q), GST-Ku70(539-556,K539R/K542R/K553R/K556R)-GFP(Ku70NLSK539R/K542R/K553R/K556R), GST-NLSc-GFP (SV40 TNLS), or GST-GFP (Empty) were microinjected into the cytoplasm of HeLa cells. After incubation at 37°C for 30 min, the cells were fixed, and the localization of microinjected proteins was examined by fluorescence microscopy. **B** Effect of amino acid substitutions in the Ku70 NLS on binding to Impα. GST pull-down assays were performed as described in the text. The binding activity of GST-Ku70(539-556)-GFP and its mutant proteins to Impα in the presence of Impβ was analyzed by probing immunoblots with antibodies against Impα.

Our previous findings demonstrated that transport of the Ku70 NLS to the nuclear rim is mediated by the two components of a nuclear pore-targeting complex, Impα/Impβ (Koike et al, 1999c). To determine whether acetylation of the Ku70 NLS affects the interaction between Ku70 and the Impα/Impβ complex, a pull-down assay was performed. The Ku70-Impα interaction was evaluated in the presence of Impβ. As shown in Figure 4B, both the wild-type Ku70 NLS and NLS of the SV40 T antigen (positive control) efficiently bound to Impα, whereas the Ku70 double mutant (K553Q/K556Q), as well as the Ku70 quadruple mutant (K539Q/K542Q/K553Q/K556Q), exhibited reduced interaction with Impα.

In our previous study, the Impα or Impβ monomers were incapable of transporting Ku70 to the nuclear rim (Koike et al, 1999c). Taken together, these data suggest that the loss of nuclear accumulation of the Ku70 double mutant (K553Q/K556Q) is due to decreased interaction with the Impα/Impβ complex and that the acetylation of two specific Ku70 lysine residues, K553 and K556, modulates nuclear import.

### Estimating the effect of acetylation of lysine residues in the Ku70 NLS using molecular dynamics simulations

In the present study, mutants mimicking acetylation and deacetylation (KQ and KR, respectively) were utilized to evaluate the effect of acetylation at lysine residues in the Ku70 NLS. However, the properties of KQ or KR substitutions do not properly represent the acetylated state of lysine residues, and it is not easy to analyze the effects of acetylation experimentally due to the difficulty of independently controlling the acetylation of single lysine residues.

To theoretically estimate the binding energy between the Ku70 NLS and Impα, molecular models of the full-length wild-type or mutant Ku70 NLS in complex with Impα were constructed based on the reported crystallographic structure, and 5-ns molecular dynamics (MD) simulations were then carried out. In each simulation, the root mean square deviation (RMSD) values of α-carbons in the interaction site of Impα (from Q70 to N346) increased sharply during the first 100 ps and then fluctuated around their respective averages (Figure EV2), although structures of the C-terminal region of Impα and the region except the interaction site in the Ku70 NLS varied dynamically (Figure EV3). The RMSD values of all of the simulated molecules after 3 ns of the MD simulation suggested that the simulated molecules reached stable conditions with few structural changes occurring.

Distributions of the total binding energy (□*G*_*btot*_) and total gas phase energy (□*G_gas_*) between the Ku70 NLS and Impα calculated from the latter 2 ns of 5-ns MD simulation data are summarized in Figure 5. The □*G_gas_* term principally consists of electrostatic energy, excluding the contribution of solvation effects of the simulated system, and thus, it is strongly influenced by the total charge of amino acids in the contact region between the Ku70 NLS and Impα. In all three trials, the total gas phase energy between the Ku70 NLS and Impα was markedly suppressed when all lysine residues of interest were substituted with glutamine (K→Q in Figure 5B) or alanine (K→A in Figure EV4B) residues, whereas they remained at the same level as that of wild type when lysine residues were substituted with arginine residues (K→R in Figure 5B).

**Figure 5.**
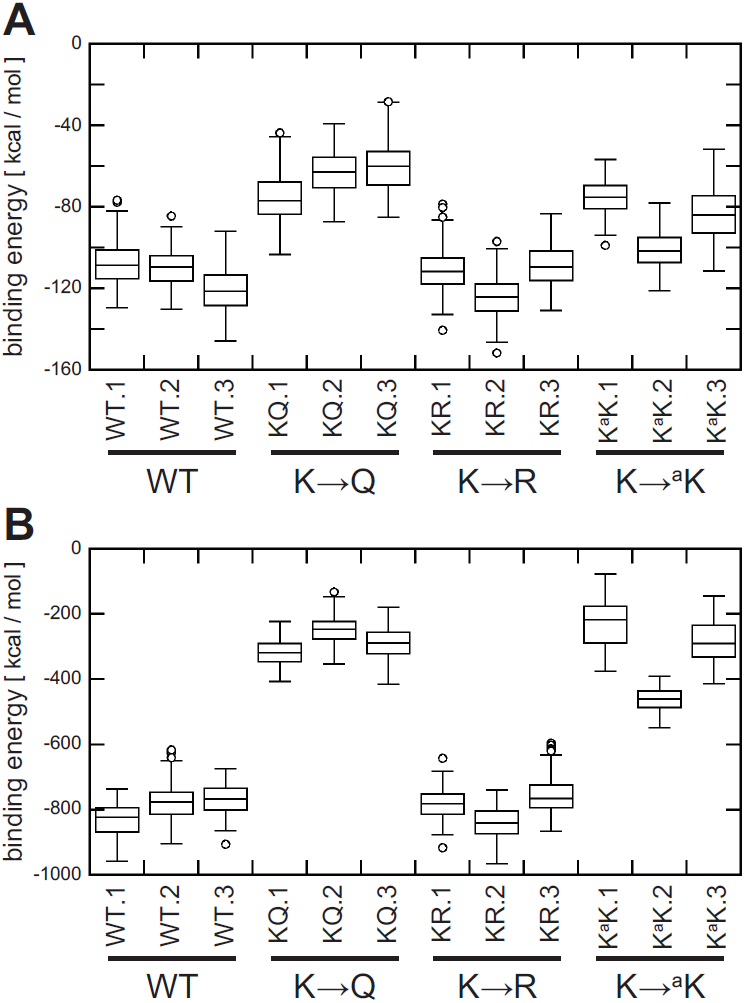
Box plots of the distribution of binding energy between Impα and the Ku70 NLS during the simulation time. **A, B** Total binding energy (□*G*_*btot*_) in solution and total gas phase energy (□*G_gas_*) between Impα andwild-type Ku70 NLS (WT.1-WT.3), KQ mutants (KQ.1-KQ.3), KR mutants (KR.1-KR.3), or K^a^K mutants (K^a^K.1-K^a^K.3) of Ku70 NLS. Data information: Each plot represents the distribution computed using coordinate sets resulting from three independent trials of 5-ns MD simulations; coordinate data were sampled every 10 ps from the latter 3 ns of each simulation. Note that lower binding energies indicate stronger interactions. Boxes represent half the number of data in each respective distribution. The bars in each box indicate the median of the distribution. Open circles represent outlying data.

This order of binding energy in the gas phase was maintained even when solvation effects were considered (Figure 5A, Figure EV4A). Less interaction between Ku70 NLS KQ mutants and Impα compared with the wild-type Ku70 NLS or its KR mutants was considered to impede binding between Impα and Ku70, which inhibits translocation of Ku70 to the nucleus. The experimental results described above strongly support this conclusion. When the lysine residues of interest were replaced with acetyl lysine residues, the binding energy in the gas phase decreased to the same level as that of KQ substitutions (K→^a^K in Figure 5B), and, in solution, approximately to the middle level between the wild type and KR substitution mutants (K→^a^K in Figure 5A), although the distribution in each trial varied over a wider range than that of the wild type or mutants.

To verify the stability of the Ku70 NLS–Impα complex, distances from a conserved residue at the contact site in the Ku70 NLS to adjacent residues in Impα during the simulation time from 3 to 5 ns were plotted (Figure 6). Distances between K553, R554, and K556 in the wild-type Ku70 NLS and their adjacent residues in Impα and between R553, R554, and R556 in the KR mutant and the same residues in Impα remained within approximately 2∼3 Å during the simulation time. In contrast, distances between residues in Impα and the KQ mutant varied and did not remain constant. When K553 and K556 were replaced with acetylated lysine residues, the distances ^a^K553 to N188, ^a^K553 to D192, R554 to N228, and R554 to D270 varied as in the case of the KQ mutant, but the ^a^K556 to W142 and ^a^K556 to W184 distances remained the same as in the wild type or KR mutant, indicating that ^a^K556 continues to interact with W142 and W184 in Impα, and thus, this could be the reason why the Ku70 NLS with a K^a^K substitution would have a slightly stronger binding energy with Impα than the KQ mutant (K→^a^K in Figure 5A).

**Figure 6.**
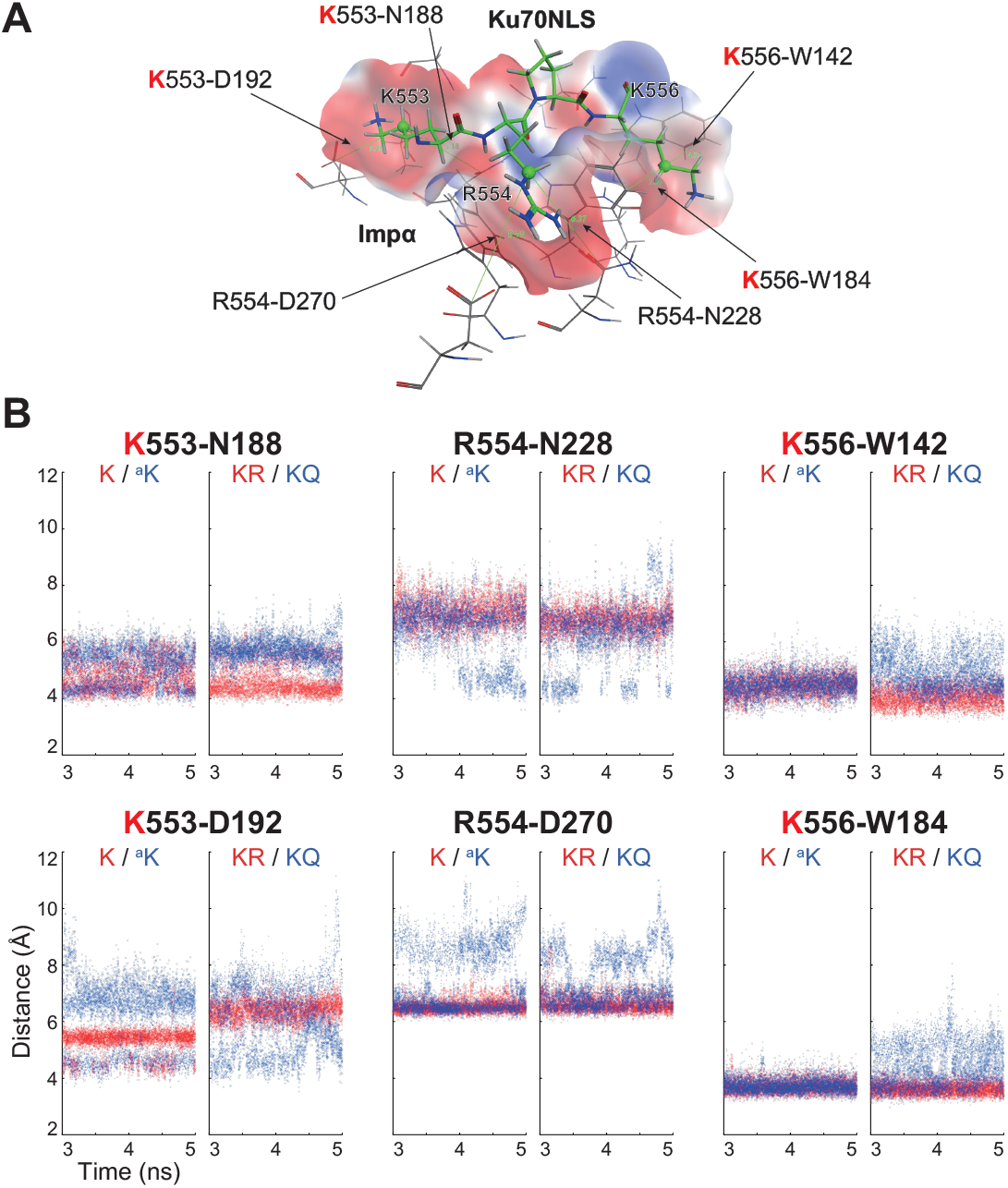
Distances from a conserved reside in the contact site of the Ku70 NLS to adjacent residues in Impα during the simulation time from 3 to 5 ns. **A** Snapshot of the interaction site between Ku70 NLS and Impα. ^553^KRPK^556^ of Ku70 NLS is illustrated as a green stick, and amino acids of Impα located near four Ku70 residues are presented as a gray stick. The molecular surface of Impα near the four Ku70 residues is also shown, with blue or red color on the surface indicating positive or negative charge, respectively. **B** Distances between CD atom in K553 of Ku70 and CG atoms in N188 and D192 of Impα, CD atom in R554 of Ku70 and CG atoms in N228 and D270 of Impα, and CD atom in K556 and CZ3 atoms in W142 and W184 of Impα are plotted during the simulation time from 3 to 5 ns (CD atoms in K553, R554, and K556 of the Ku70 NLS are represented as green balls in **A**. In each panel, the left graph represents the distribution of distances when none of the acetylation targets in the Ku70 NLS were replaced (K) or when they were replaced with acetyl lysine (^a^K) residues, and the right graph indicates the distribution of distances when all acetylation targets in the Ku70 NLS were replaced with arginine (R) or glutamine (Q) residues.

Taken together, the results of the theoretical analysis strongly suggest that interaction between the Ku70 NLS and Impα is impeded when lysine residues in the Ku70 NLS are acetylated, which inhibits the ability of Impα to transfer Ku70 to the nucleus.

## Discussion

Spatiotemporal regulation of PTMs for DNA repair proteins responding to DNA damage is essential for the precise targeting and control in the various DNA repair pathways (Koike, 2002; Koike et al, 2017a). Accurate PTM-mediated control of Ku70 subcellular localization, recruitment to DSBs, and protein-protein interactions might play a central role in the regulation of not only NHEJ activity but also multiple Ku70-dependent biological processes, including telomere maintenance and apoptosis (Koike, 2002; Koike et al, 2017a). It was reported that there are at least eight acetylation target sites in Ku70 (Cohen et al, 2004a). Ku70 is acetylated by acetyltransferases such as CBP and PCAF, whereas it is deacetylated by various histone deacetylases, including HDAC3, HDAC6, SIRT1, and SIRT6 (Cohen et al, 2004a; Cohen et al, 2004b; Gong et al, 2018; Subramanian et al, 2011; Tao et al, 2017). These data strongly suggest that the multiple functions of Ku70 are, at least in part, regulated by its acetylation, although the biological significance of acetylation of Ku70 has not been thoroughly elucidated.

To evaluate the effects of acetylation experimentally, recombinant mutants in which lysine residues were substituted with glutamine residues to mimic acetylated lysine or with arginine to mimic deacetylated lysine, preserving the positive charge from acetylation, have been employed for a wide variety of proteins (Kim et al, 2006), including histones H3 and H4 (Hecht et al, 1995; Johnson et al, 1990) and p53 (Li et al, 2002). However, the properties of these substitutions have not been properly validated due to the difficulty of independently controlling acetylation of single lysine residues, though a few alternative experimental methods have been developed (Cohen et al, 2004a; Kamieniarz & Schneider, 2009; Li et al, 2002; Neumann et al, 2008).

We previously examined the effect of acetylation of three lysine residues inside the Ku70 ring, K317, K331, and K338, on the binding affinity of Ku for DNA using the same molecular simulation technique described in this report and demonstrated that acetylation of these lysine residues in the Ku70 ring does not markedly reduce the affinity of Ku for DNA, although those residues have been acetylated in *in vivo* studies (Cohen et al, 2004a) and the binding affinity of Ku for DNA was decreased *in vitro* when those residues were substituted with glutamine residues (Chen et al, 2007). The evidence that EGFP-Ku70 K338Q in the presence of Ku80 is capable of accumulating at DSB sites in living cells partly supports the result of the theoretical analysis (Koike et al, 2011). These observations also suggest that the effect of *in vivo* acetylation is sometimes overestimated when the KQ mutant is employed as a mimic of the acetylated protein (Fujimoto et al, 2012). In the present study, the level of nuclear localization of Ku70 was found to decline when some of the lysine residues in the Ku70 NLS were replaced with arginine residues (e.g. ---R-, ---RR in Figure 2D). In particular, the ability to translocate Ku70 to the nucleus was almost completely abolished when all five lysine residues were substituted with arginine residues (RRRRR in Figure 2D). This finding indicates that, for translocation of Ku70 to the nucleus, it is important that all or some of these positions are occupied by lysine residues, and translocation cannot proceed with substituting all of the lysine residues for arginine residues. This suggests that similar to the case of the KQ mutant, the KR mutant does not exactly mimic the deacetylated protein.

The molecular structures of the core of the Ku heterodimer and C-terminal DNA-binding domain (SAP domain) of Ku70 were separately resolved by X-ray crystallographic and NMR analysis, respectively (Walker et al, 2001; Zhang et al, 2001). The Ku70 NLS is located in the linker region connecting these two portions of Ku70 and is thought to be incapable of folding by itself into a specific three-dimensional (3D) structure. This disordered region is probably flexible in solution and has enough length for regulatory proteins, such as Impα, to associate with Ku70 without sterically interfering with the rest of the Ku70 molecule (Figure EV5).

In this study, the function of acetylation and deacetylation of five lysine residues, K539, K542, K544, K553, and K556, in the Ku70 NLS was analyzed using both experimental and theoretical approaches. Two of these five lysine residues in the C-terminus of the Ku70 NLS, K553 and K556, were identified as key players in modulating the nuclear transport of Ku70. Of the remaining three lysine residues, K539 and K542 were found to contribute less to both the dimerization of Ku70 and Ku80 and to the nuclear transport function, because mimicking the acetylation of either or both of these lysine residues had minimal impact on nuclear transport (Figure 2), and EGFP-Ku70 K539Q, EGFP-Ku70 K542Q, and EGFP-Ku70 K539Q/K542Q maintained the ability to accumulate at DSB sites in living cells (Figure 3B). It was reported that the acetylation of Ku70 regulates two independent apoptosis pathways (i.e., Bax-mediated apoptosis and Bax-unmediated FLIP/caspase-8–dependent apoptosis) (Cohen et al, 2004a; Kerr et al, 2012). Interestingly, the substitution of either K539 or K542 with glutamine abolished the binding of cytosolic Ku70 to both Bax and FLIP (Cohen et al, 2004a; Kerr et al, 2012; Subramanian et al, 2005). Therefore, the biological consequences of acetylation of the five lysine residues in the Ku70 NLS appear to be bifunctional; acetylation of K553 and K556 suppresses the nuclear transport of Ku70 for DSB repair, whereas acetylation of K539 or K542 facilitates the activation of the two apoptosis pathways, although the biological significance of the acetylation of K544 in the Ku70 NLS remains unclear.

Ku80 also has a region capable of interacting with Impα in the linker region connecting the Ku80 core with the C-terminal protein-binding domain (Koike et al, 1999b; Takeda et al, 2011), suggesting that the nuclear transport processes involving Ku70 and Ku80 are regulated independently, although it remains unclear whether the translocation of Ku80 to the nucleus is controlled by acetylation or deacetylation of lysine residues in the Ku80 linker region. Surprisingly, some other components associated with the NHEJ pathway, such as XRCC4, have a similar NLS in which the lysine residues are conserved (Koike et al, 2017b). Further structural analyses of NHEJ enzymes are required.

Our findings strongly suggest that the acetylation of certain lysine residues in the Ku70 NLS is a key switch that controls the localization of Ku70 by modulating its interaction with nuclear transport factors. Nuclear Ku70 is an essential factor in DNA repair pathways, NHEJ, and telomere maintenance, and cytosolic Ku70 might be an apoptosis regulator. These functional distinctions raise the question as to whether Ku70 plays a role in determining the destiny (i.e., life or death) of cells with damaged DNA. Moreover, various reports recently showed that Ku70 can function in the cytoplasm with or without binding to Ku80 with respect to virus infections. For instance, as a cytosolic DNA sensor, Ku70 translocates from the nucleus to the cytoplasm and forms a complex with STING, which regulates the Ku70-mediated production of IFN-λ1 in response to exogenous DNA, such as DNA derived from HSV-2 infection (Sui et al, 2017). In sensing HBV DNA, the sensing of HBV DNA by the cytoplasm-localized Ku70/Ku80 complex induces hepatitis-associated chemokine secretion (Li et al, 2016). These data along with the findings of the present study indicate that precise spatiotemporal regulation of PTMs in Ku70 is a crucial factor not only with respect to the DNA damage response but also responses to virus infections. The subcellular localization and PTMs of Ku70 change dynamically during the cell cycle (Koike et al, 1999a; Koike et al, 2017a; Mahaney et al, 2009). Further experimental and theoretical studies, including molecular structural analyses of acetylation of Ku70, might be useful for clarifying the mechanism by which the multiple functions of Ku70 are regulated in both the nucleus and cytoplasm. The combined analytical method described here provides a novel approach for analyzing the control mechanism for other acetylated proteins.

## Materials and Methods

### Cell lines, culture, and transfection

As described previously, human cervical carcinoma cells (HeLa) were cultured in Dulbecco’s modified Eagle’s medium supplemented with 10 % fetal bovine serum (FBS) and antibiotics. Hamster xrs-6 cells defective in Ku80 (derived from CHO-K1 cells on the basis of their sensitivity to ionizing radiation) (Koike & Koike, 2004; Singleton et al, 1997) were cultured in Ham’s F12 medium (Sigma, St. Louis, MO, USA) supplemented with 10% FBS and antibiotics. The xrs-6 cells were plated in a 35-mm dish (Falcon, Corning, NY, USA) at a density of 1.5 ∼ 2.0 × 10^5^ cells/well the night before transfection. Cells were transfected using FuGene6 (Roche Diagnostics, Indianapolis, IN, USA) or Lipofectamine 3000 (Invitrogen, Carlsbad, CA, USA).

### Plasmid construction

cDNAs for human Ku70 and Ku80 were derived from pEGFP-Ku70(1-609) or pECFP-Ku80(1-732) (Koike et al, 1999c; Koike et al, 2001). Ku70 site-specific mutants (pEGFP-Ku70 K539Q, pEGFP-Ku70 K542Q, pEGFP-Ku70 K544Q, pEGFP-Ku70 K553Q, pEGFP-Ku70 K556Q, pEGFP-Ku70 K539R, pEGFP-Ku70 K542R, pEGFP-Ku70 K544R, pEGFP-Ku70 K553R, pEGFP-Ku70 K556R, pEGFP-Ku70 K539Q/K542Q, pEGFP-Ku70 K553Q/K556Q, pEGFP-Ku70 K539R/K542R, pEGFP-Ku70 K553R/K556R, pEGFP-Ku70 K539A/K542A, pEGFP-Ku70 K553A/K556A, pEGFP-Ku70 K539Q/K542Q/K553Q/K556Q, pEGFP-Ku70 K539R/K542R/K553R/K556R, or pEGFP-Ku70 K539Q/K542Q/K544Q/K553Q/K556Q,pEGFP-Ku70K539R/K542R/K544R/K553R/K556R,orpEGFPKu70K539A/K542A/K544A/K553A/K556A) were generated by incorporating mutant oligonucleotides via strand extension reactions, as previously described (Koike et al, 1999b; Koike et al, 2001). Following the application of the mutagenesis strategy, each mutant was identified by DNA sequencing, as previously described (Koike et al, 1999c).

### Inducing local DNA damage using laser irradiation and cell imaging

As described previously, local DNA damage was induced using a laser, and cells were subsequently imaged (Koike & Koike, 2008; Koike et al, 2011). Briefly, local DSBs were induced using a 10% power scan (for 1 s) from a 405-nm laser. Images of living or fixed cells expressing EGFP-tagged proteins or EGFP alone were acquired using an FV300 CLSM system (Olympus Co., Tokyo, Japan).

### Preparation of GST-Ku70-GFP fusion proteins

To construct GST-Ku70-GFP fusion genes, the GST-GFP2 cassette vector (Eguchi et al, 1997) was used. The sequences of the oligonucleotides used to prepare fragments of Ku70 were as follows: 404 (<u>GATCCCT</u> AAA GTT ACC AAG AGA AAA CAC GAT AAT GAA GGT TCT GGA AGC AAA AGG CCC AAG G) and 405 (<u>GATCCCT</u> TGG GCC TTT TGC TTC CAG AAC CTT CAT TAT CGT GTT TTC TCT TGG TAA CTT TAG G) for the wild-type Ku70 NLS (539-556); SMF80 (<u>GATCCCT</u> AAA GTT ACC AAG AGA AAA CAC GAT AAT GAA GGT TCT GGA AGC CAA AGG CCC CAG G) and SMR80 (<u>GATCCCT</u> GGG GCC TTT GGC TTC CAG AAC CTT CAT TAT CGT GTT TTC TCT TGG TAA CTT TAG G) for Ku70 NLS (539-556)/(K553Q/K556Q); 398 (<u>GATCCCT</u> CAA GTT ACC CAG AGA AAA CAC GAT AAT GAA GGT TCT GGA AGC CAA AGG CCC CAG G) and 406 (<u>GATCCCT</u> GGG GCC TTT GGC TTC CAG AAC CTT CAT TAT CGT GTT TTC TCT GGG TAA CTT GAG G) for Ku70 NLS (539-556)/(K539Q/K542Q/K553Q/K556Q); and 400 (<u>GATCCCT</u> AGA GTT ACC AGG AGA AAA CAC GAT AAT GAA GGT TCT GGA AGC AGA AGG CCC AGG G) and 407 (<u>GATCCCT</u> GGG CCT TCT GCT TCC AGA ACC TTC ATT ATC GTG TTT TCT CCT GGT AAC TCT AGG) for Ku70 NLS (539-556)/(K539R/K542R/K553R/K556R). Each pair of oligonucleotides was annealed, phosphorylated, and then ligated to GST-GFP2 at the *Bam*HI site. The GST-NLSc-GFP construct, which contained the core sequence of the NLS of the SV40 T-antigen, was prepared as described previously (Eguchi et al, 1997). The GST-Ku70-GFP vectors described above were introduced into *Escherichia coli* strain BL21. Purification of expressed GST-Ku70-GFP fusion proteins was carried out as described previously (Eguchi et al, 1997). The purified proteins were dialyzed against PBS.

### Microinjection into the cytosol of HeLa cells

Microinjection experiments were performed as described previously (Eguchi et al, 1997; Koike et al, 1999b; Yoneda et al, 1987). Various GST-Ku70-GFP fusion proteins were injected into the cytosol of HeLa cells via microinjection, and the cells were incubated at 37°C for 30 min before fixation with 4% formaldehyde. Fusion proteins were localized using fluorescence microscopy.

### Binding assay

GST-Ku70-GFP proteins, adjusted to the same amount by comparison with the intensity of SDS-PAGE bands, were incubated with glutathione Sepharose 4B (Thermo Fisher Scientific Inc. (Invitrogen), Waltham MA, USA) for 1 h at 4°C and then washed three times with transport buffer (pH 7.3), consisting of 20 mM Hepes, 110 mM potassium acetate, 2 mM magnesium acetate, 5 mM sodium acetate, 1 mM EDTA, 1 mM EGTA, and 2 mM dithiothreitol. The Sepharose was incubated with purified PTAC58 (Impα) and PTAC97 (Impβ) in transport buffer containing 1 mg/ml BSA. After incubation for 1 h at 4°C, the Sepharose was washed three times with transport buffer, added to lysis buffer, and boiled for 5 min. The eluted proteins were separated by 8% SDS-PAGE and blotted onto a nitrocellulose membrane. The membrane filter was incubated with an Impα-specific antibody, anti-KPNA2 (Everest Biotech Ltd., Oxfordshire, UK), and then incubated with alkaline phosphatase–conjugated anti-goat IgG. After washing with buffer, the bound antibodies were detected using an alkaline phosphatase conjugate substrate kit (Bio-Rad, Hercules, CA, USA).

### Modeling of the Ku70 NLS–Impα complex

A molecular model of the Ku70 NLS–Impα complex was constructed based on the reported crystallographic structure (PDB ID: 3RZX) (Takeda et al, 2011). Missing atoms in the structural data for Impα were complemented using the 2016.0802 version of the Molecular Operating Environment (MOE) (Chemical Computing Group Inc., ontreal, Canada http://www.chemcomp.com/MOE-Molecular_Operating_Environment.htm). As the PDB data contain a partial NLS of Ku70, missing amino acid residues (^537^EGKVTKRKHD^546^) were added to the N-terminus of ^547^NEGSGSKRPKVE^558^. The five lysine residues (K) in the wild-type Ku70 NLS at positions 539, 542, 544, 553, and 556 were replaced with alanine (A), arginine (R), glutamine (Q), and acetyl lysine (^a^K) residues to prepare experimentally verifiable models. The molecular model of acetyl lysine (Nε-acetyl-L-lysine) was established from a standard lysine molecular model. Partial charges of each atom were calculated previously (Fujimoto et al, 2012). The protonation state of each molecular model was determined using the Protonate 3D application in the MOE program package.

### MD simulation

Classical MD calculations for the Ku70 NLS–Impα complex were carried out using the SANDER module in the AMBER 14 program package (Case et al, 2014) (http://ambermd.org/). The AMBER force field, ff14SB, was applied to all simulated systems. The negative charge of each system was neutralized by the addition of sodium counter ions (Na^+^), and each modeled complex was immersed in solute molecules in a rectangular parallelepiped filled with a periodic box of TIP3P water molecules (Jorgensen et al, 1983). The potential energy of each solvated system was minimized to optimize the initial position of each atom. Details of the MD protocol were described previously (Fujimoto et al, 2012). Through all simulations, time steps were defined as 1 fs, and the cutoff distance of the van der Waals interactions was set at a 10 Å. After the minimization step, the simulated system was heated from 0 to 310 K (∼37°C) for 10,000 steps (10 ps) by applying kinetic energy to each atom, and then the density of the system was saturated at a constant pressure of 1 atm for 10,000 steps (10 ps). Productive MD calculations were performed in a constant rectangular volume for up to 5,000,000 steps (5 ns). Long-range electrostatic interaction energies were calculated using the particle-mesh Ewald method (Darden et al, 1993; Essmann et al, 1995), and the SHAKE algorithm (Andersen, 1983; Ryckaert et al, 1977; van Gunsteren & Berendsen, 1977) was applied to constrain all bonds involving hydrogen atoms during the heating and constant-pressure steps. The coordinate sets and energies of each atom were stored at 1-ps intervals for further analysis. To avoid locally stable conditions, the process was repeated three times for each model, with different initial velocities set for all atoms in the system before the start of the heating step.

### Estimation of binding free energy

Binding free energies between the Ku70 NLS and Impα were calculated according to the MM/PBSA method (Kollman, 1993) based on the following equation: □*G_btot_* = □*G_gas_* + □*G_sol_* □*T*□*S*, where □*G*_*btot*_ represents the binding free energy in solution, □*G_gas_* represents the total gas phase energy that was, in this study, equal to the sum of the electrostatic energy and the van der Waals interaction energy between the Ku70 NLS and Impα in the gas phase, and □*G*_*sol*_ represents the solvation energy. □*T*□*S* represents the contribution of entropy, but this term was omitted in this study because the differences between the entropy contribution of the wild-type Ku70 NLS-Impα complex and that of various mutants was assumed to be negligible. Every term, with the exception of □*T*□*S*, was calculated using the mm_pbsa.pl module in the AMBER 14 software package using default parameters for 200 coordinate sets selected from the latter 2 ns of a 5-ns MD simulation every 10 ps. The Poisson-Boltzmann model was applied as a solvent model to calculate solvation energy.

## Acknowledgements

We thank Dr. P. Jeggo for providing xrs-6 cells and Dr. H. Ode for preparing the computer script to assemble MM/PBSA data.

## Author contributions

Fujimoto H developed the concept and design of the theoretical portion of this study and performed molecular simulations. Ikuta T performed the pull-down and *in vitro* transport assays and experimentally evaluated Ku70-Impα interactions. Koike A and Koike M carried out live-cell imaging and assembled the data. Koike M developed the concept and design of the experimental portion of this study. Fujimoto H and Koike M wrote most of the manuscript, and Koike A and Ikuta T read and approved the final version of the manuscript.

